# Association Between Polygenic Risk Score And Gut Microbiome Of Multiple Sclerosis

**DOI:** 10.1101/2022.11.07.515482

**Authors:** Noha S. Elsayed, Robert K. Valenzuela, Terrie Kitchner, Thao Le, John Mayer, Zheng-Zheng Tang, Vishnu R. Bayanagari, Qiongshi Lu, Paula Aston, Karthik Anantharaman, Sanjay K. Shukla

**Author notes:** **Corresponding author** Sanjay K. Shukla, PhD, Center for Precision Medicine Research, Marshfield Clinic Research Institute, 1000 N Oak Avenue # MLR, Marshfield, WI 54449. Roger Williams Medical Center - Boston University School of Medicine Providence, RI, 02908. **Emails:** Noha S.Elsayed Robert K. Valenzuela Terrie Kitchner, Thao Le John Mayer Zheng-Zheng Tang Vishnu R. Bayanagari Qiongshi Lu Paula Aston Karthik Anantharaman Sanjay K. Shukla.

## Abstract

**Background:** Multiple sclerosis (MS) is a complex autoimmune disease in which both the roles of genetic susceptibility and environmental/microbial factors have been investigated. More than 200 genetic susceptibility variants have been identified along with the dysbiosis of gut microbiota, both independently have been shown to be associated with MS. We hypothesize that MS patients harboring genetic susceptibility variants along with gut microbiome dysbiosis are at a greater risk of exhibiting the disease. We investigated the polygenic risk score for MS in conjunction with gut microbiota in the same cohort of 117 relapsing remitting MS (RRMS) and 26 healthy controls. DNA samples were genotyped using Illumina’s Infinium Immuno array-24 v2 chip followed by calculating polygenic risk score and the microbiota was determined by sequencing the V4 hypervariable region of the 16S rRNA gene.

**Results:** We identified two clusters of MS patients, Cluster A and B both having a higher polygenic risk score than the control group. The Cluster B with the higher polygenic risk score had a distinct gut microbiota, different than the Cluster A. MS group whose microbiome was similar to that of the control group despite a higher genetic risk score than the control group. This could be due to i) the non-active state of the disease in that group of MS patients at the time of fecal sample collection and/or ii) the restoration of the gut microbiome post disease modifying therapy to treat the MS.

**Conclusion:** Our study showed that there seems to be association between polygenic risk score and gut microbiome dysbiosis in triggering the disease in a small cohort of MS patients. The MS Cluster A who have a higher polygenic risk score but microbiome profile similar to that of healthy controls could be due to the remitting phase of the disease or due to the effect of disease modifying therapies.

## Background

Multiple sclerosis (MS), a disease that affects nearly 2.8 million people worldwide [1], is a chronic, inflammatory, autoimmune disease of the central nervous system with a complex, multifactorial etiology [2]. The symptoms of MS range from fatigue, numbness, muscle spasms and weakness to various gastrointestinal and urinary malfunction symptoms [3]. Pathologically, the disease manifests with demyelination and degeneration of neurons, and presence of white matter lesions on the brain and the spinal cord [1,3]. What etiological factors drives the two phenotypes of MS: relapsing-remitting MS (RRMS) and primary progressive MS (PPMS) [3] is not fully understood. The most common phenotype is the RRMS where the patient alternates between active and non-active episodes of symptoms. The active episodes are marked with motor, sensory and cognitive symptoms in addition to brain lesions detected by magnetic resonance imaging [4]. The complex etiology of MS disease continues to being investigated through increasing understanding of genetic susceptibility and different triggering modalities arising from life-style and/or environment [5] such as smoking, low sun light exposure, high salt diet, viral infection(s), and microbe(s) or microbial metabolites emanating from gut microbiome dysbiosis [6–9]. The gut microbiome with its dynamic reservoir of trillions of microbes representing hundreds of species is of great interest to potentially link its role in genetically susceptible persons.

Indeed, genetic susceptibility to MS is complex and hundreds of genomic regions that are implicated are dispersed throughout the chromosomes [10]. The genetic susceptibility accounts for 30% of the MS cases [11]. Siblings of MS patients are seven times more susceptible for this disease than general population [12]. However, the major histocompatibility complex (MHC) haplotypes on chromosome 6 have shown as the highest reproducible associations with MS susceptibility. Mostly MHC class II alleles, such as DQA1*01:01-DRB1*15:01 and DQB1*03:01-DQB1*03:02 are pivotal [13]. The strongest risk allele is HLA-DRB1*15:01 with an odds ratio of 3.08 [14]. Usually, in complex diseases like MS, the more risk alleles the subject carries, the higher the predisposition to the disease [15]. Thus, measuring the polygenic risk score of the MS patients in comparison to the healthy controls can be used in revealing more precise genetic susceptibility to this complex disease.

While knowledge of genetic susceptibility to MS has enhanced our understanding of the disease, the precise source and role of the environmental factor(s) including microbial trigger(s) associated with MS is far from settled. The human gut microbiome with its rich source of microbial diversity, their antigens, and metabolites are being explored as a possible source of infectious triggers modulating the MS disease. Indeed, several recent studies have reported association of gut microbiome dysbiosis with the MS [2,6,16]. A convincing role for the gut microbiome in the MS disease was supported by observation from an experimental autoimmune encephalomyelitis (EAE) disease mouse model analogous to MS. In this model, SJL/J mice were protected from MS when grown in germ free conditions [17]. Furthermore, their susceptibility to EAE was restored by exposing these mice to the commensal bacteria from fecal material from specific pathogen-free mice [17]. Additionally, the MS disease development was reproducible in a EAE mice model by transferring MS patient’s fecal material to mice [5]. Since then, several studies have reported an association of MS with the gut microbiome dysbiosis involving different taxa. For examples *Akkermansia* and *Methanobrevibacter* are in higher relative abundance whereas *Prevotella* was in lower relative abundance [6,16,18]. However, there seems to be discrepancies in different study results with respect to experimental details and statistical analysis [19], in genetic and environmental dissimilarity [11] among MS patient cohorts or even disease treatment regimens [20].

A role of host genetics selecting and/or modulating gut microbiome in both healthy and diseased cohorts have been described particularly in type 1 diabetes and rheumatoid arthritis [21,22]. Polygenic risk score enhances the predictive power of disease susceptibility and outcome [23]. A population with both genetic and environmental risk factors (GxE) are at a greater disease risk [24]. In this study, we show that a cohort of MS patients have enhanced polygenic risk score and also harbor a distinct gut microbiota which is different from the healthy controls suggesting an association between the genetic risk score and gut microbiota.

## Methods

### Study approval

This study obtained approval from institutional research board (IRB) of Marshfield Clinic Health System under IRB protocol SHU10417 and all of the included subjects signed a written informed consent. The reporting of this study followed most of the STORMS checklist for microbiome reporting studies [25].

### Study design

Two-hundred thirty seven MS patients and 50 controls were recruited in this case-control study from the Marshfield Clinic-Marshfield Center during 2018-2021 who have had a recent diagnosis of MS (< 2 years of disease duration) or established diagnosis of MS (> 2 years of disease duration) regardless of clinical subtype (PPMS and RRMS) and treatment modality. The exclusion criteria were patients taking antibiotics, laxatives, or probiotics or who underwent a colonoscopy or similar procedure during the last three months.

All 237 patients provided a ∼ 5.0 ml of blood samples while only 214 patients provided their fecal sample. All 50-control subjects provided both blood and fecal samples. We determined the 16S-based microbiota from 169 cases and 33 controls. The 169 patients were binned into five groups: treated RRMS (Group 1), treated PPMS (Group 2), treatment naïve RRMS but diagnosed for >2 years of disease duration (Group 3), treatment naïve RRMS diagnosed for <2 years of disease duration (Group 4), and treatment naïve PPMS (Group 5) as shown in **Figure 1**. The patients included in groups one and two were on disease-modifying treatments (DMT) within six months of their stool collection. The DMTs were Glatiramer acetate, Dimethyl fumarate, Fingolimod, Natalizumab, Ocrelizumab or Teriflunomide. Groups two, four, and five were excluded from further analysis because each included <10 patients, and 36 cases from Group 1 and Group 3 and seven healthy controls samples were filtered out due to low sequencing reads. The final microbiome analysis was based on 117 MS cases and 26 control subjects.

**Figure 1.**
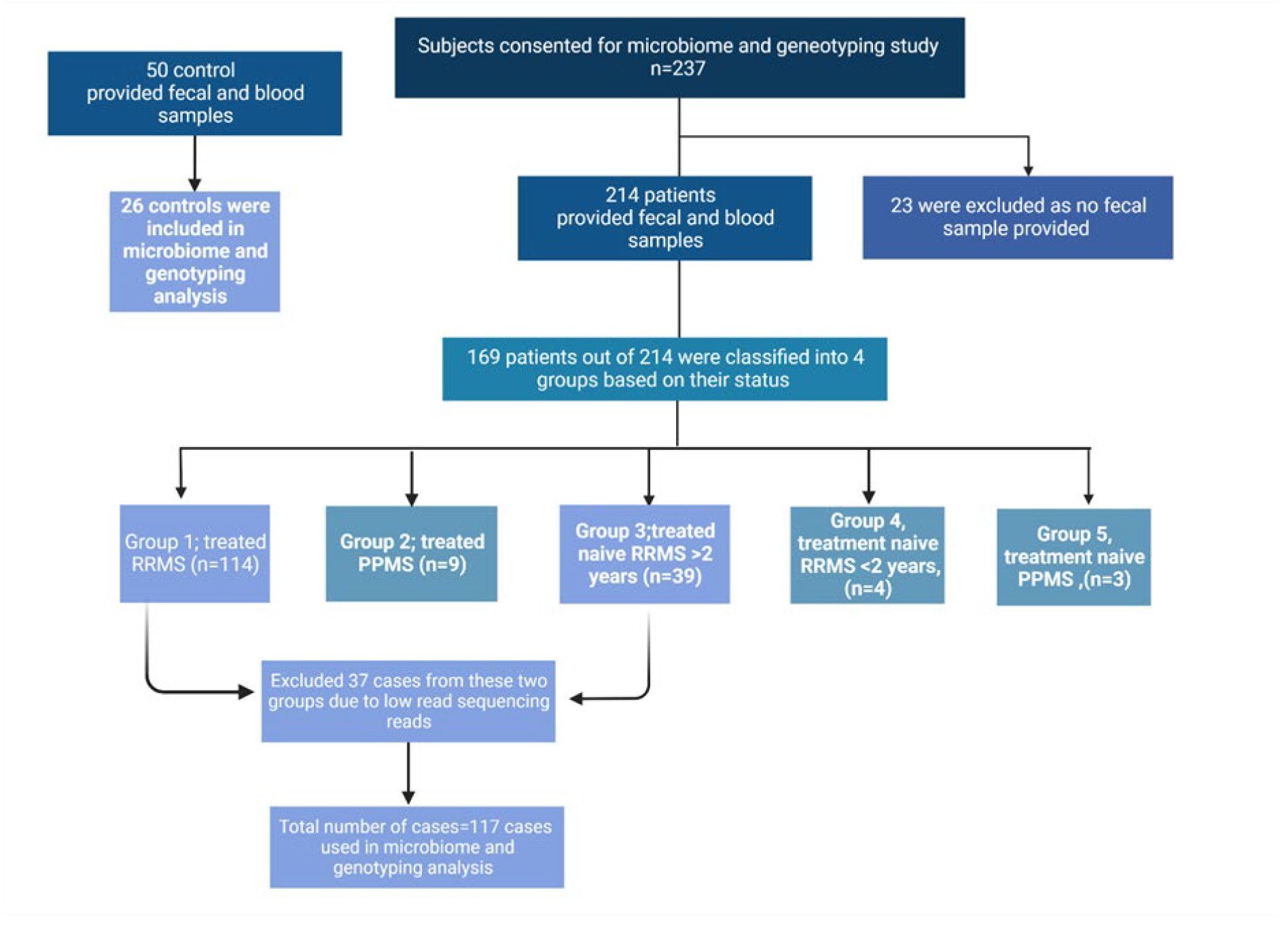
Flow chart displaying both the recruited MS patients and the healthy controls, their grouping and inclusion in the final analysis

### Sample collection and storage

A self-collection fecal sample kit with detailed instructions was sent to each subject (patients and controls) or handed over by medical assistant of the caring physician to the MS patient during their routine visit with a provider. The fecal samples were returned in a boxed frozen cold pack to Dr. Shukla’s laboratory where they were divided into aliquots and stored at −80°C until further analysis. In addition, the recruited patients and controls provided a blood sample during their regular visit to MCHS’s phlebotomy center. The blood samples were processed for serum, plasma, and buffy coat collection and stored at −80°C.

### DNA extraction and 16S rRNA amplification

The microbial DNA was extracted from the fecal material using PowerLyzer PowerSoil DNA Isolation Kit (MoBio Laboratories, Inc., Carlsbad, CA) by following the manufacturer’s protocol. Integrated DNA Technologies (Ames, IA) synthesized oligonucleotide primers (515F-806R) required for amplifying V4 region of 16S rRNA sequences [26] where the reverse amplification primer contained a 12 base barcode sequence and both primers contain adaptor regions [27]. The amplification was carried out using PE9700 thermocycler with the following run conditions initial denaturing temperature 94ºC for 2 min, 35 cycles of 94ºC for 45 seconds, 64ºC for 45 seconds and 72ºC for 45 seconds followed by a 10 min at 72ºC as final extension. SequalPrep™ Normalization Plate Kit was used to normalize the amplicon concentration (Thermofisher Scientific). Sequencing was done using the Illumina MiSeq Reagent Kit V2 with V4 sequencing primers as described by Caporaso et al (2012) [28]. The total number of reads was 8,785,102 with an average read of 43,491.

### Sequence data analysis

The demultiplexed paired-end reads from MiSeq were imported into Quantitative Insights Into Microbial Ecology (QIIME2, version 2019.10) [29] custom pipeline where the reads were assembled into one Fastq file identified with the sample names. Then, DADA2 plugin was used to denoise the sequences [30]. A fragment insertion tree using the q2-fragment-insertion plugin was created depending on alignment with the Greengenes database [31]. The generated Amplicon Sequence Variants (ASVs) from DADA2 were assigned to taxonomy using a pre-trained Naive Bayes classifier including the existing taxa in the 99% Greengenes 13_8 reference specific to the V4 hypervariable region corresponding to the primers we used [32]. A sampling depth of 24,520 reads was used to normalize the features count in each sample. The 16S microbiome analysis was performed on 117 number of cases and 26 of controls.

### Microbiome analysis

Alpha diversity was computed using the Faith’s Phylogenetic Diversity which is the sum of the branch lengths of a phylogenetic tree connecting all species in the target assemblage [33], and Pielou’s evenness index [34] and Shannon indices using the Qiime 2 pipeline. Kruskal Wallis test was used to detect any significant differences between cases and controls in different indices of alpha diversity. Principal component analysis (PCA) based on unweighed unifrac [35] was carried out while doing the permutational multivariate analysis of variance (Permanova) test to detect if there was any significant difference between the clusters formed. Graphs were plotted using the ggplot2 package of the R statistical software 3.6.0. To detect a significant taxa at the phyla, family or genera level associated within the two clusters generated from PCA analysis, the Quasi-Conditional Association Test using General Estimating Equations (QCAT-GEE) was used, including a Permutation test [36]. The QCAT-GEE composes of three tests: the zero-test, which assess presence or absence of taxa, the positive-test, which assesses differences in abundance of each taxa, and the two-test, which combines the zero and positive-tests.

### Genotyping and polygenic risk score

Genotyping was performed on all 117 cases and 26 controls. Briefly, DNA from both the patients and healthy controls’ buffy coat was isolated using QIAamp DNA blood mini kit (Qiagen Inc; Germanton, MD). The DNA samples were genotyped using Illumina’s Infinium Immuno array-24 v2 chip at UW-Madison’s Gene Expression Center (GEC). Variants were clustered and genotyped using GenomeStudio Data Analysis software 2.0 along with the chip manifest files. The SNPs were retained for imputation based on standard criteria (e.g., minimum allele frequency > 0.05; missingness < 0.01; individual genotype rate > 0.99; and Hardy-Weinberg equilibrium p-value > 1e-07)[37]. Genetic coverage was increased through imputation using genome build 38 Genotype Imputation HLA of the University of Michigan’s Imputation Server [38]. A polygenic risk score was calculated utilizing 187 relevant variants (Table 2 and Supplemental Table 1) previously identified by Patsopolous et al (2019) [10]. The polygenic risk score as defined by Chatterjee et al (2016) is the quantitative measurement of the total genetic risk of multiple susceptibility variants (common, intermediate, and rare) of the disease [23]. The calculation of the polygenic risk score for each subject was performed by summing the number of risk alleles for a given variant and multiplying the sum by the effect size obtained from Patsopolous et al, 2019. Plink software version 2.3.1 was then used to divide the score by the total number of SNPs [39].

**TABLE 1:**
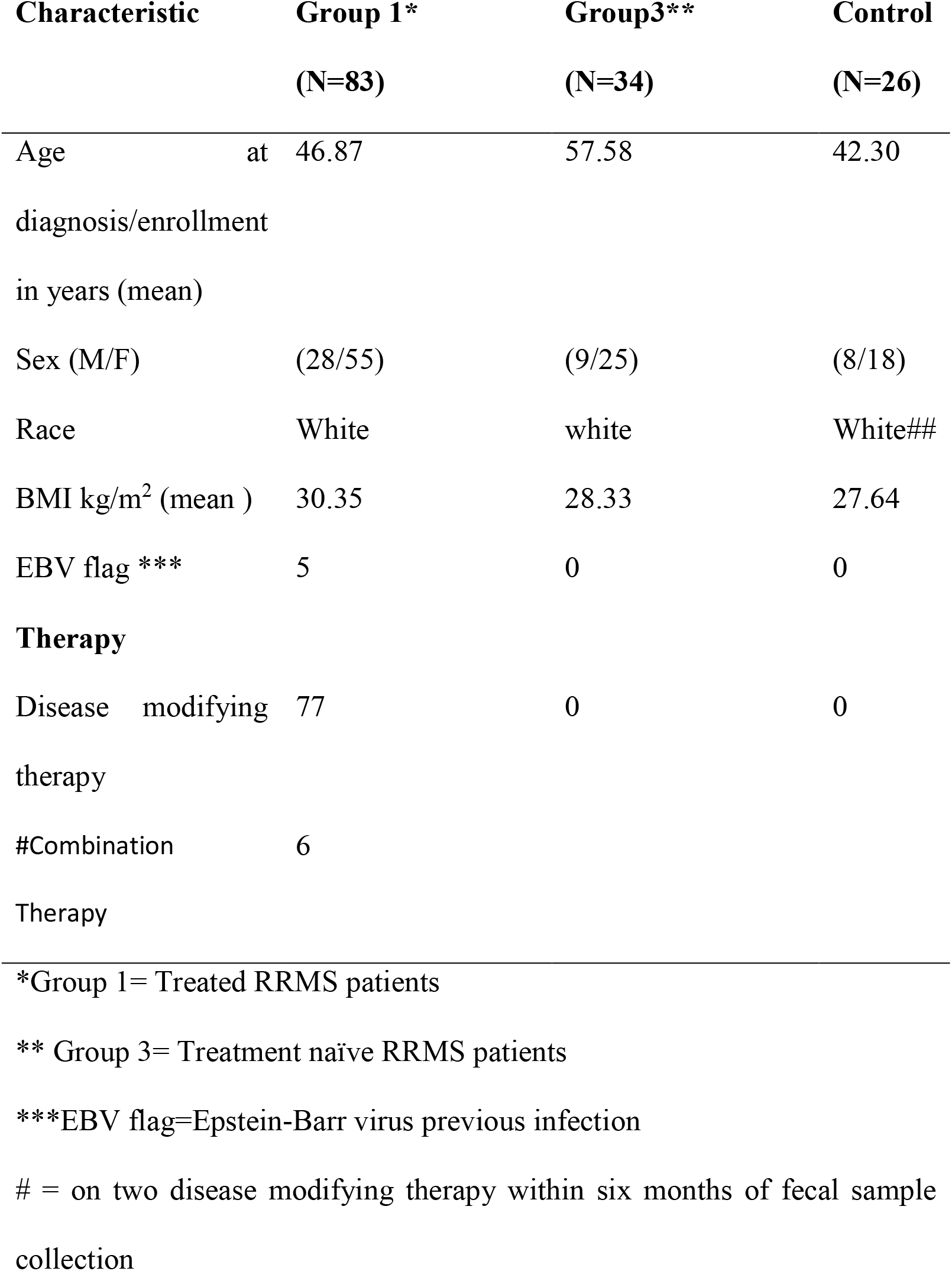
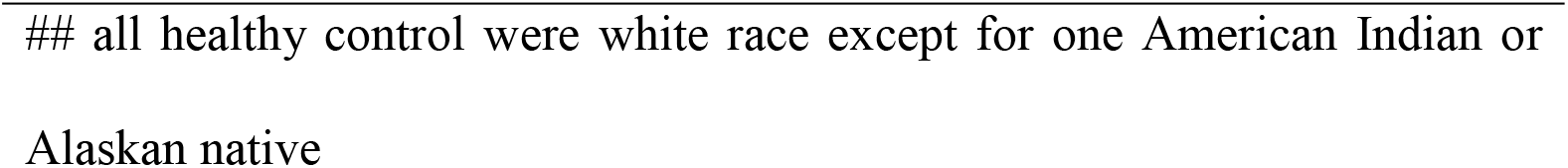
Demographic characteristics of the MS patients and healthy control at the time of the stool collection.

| Characteristic | Group 1*<br>(N=83) | Group3**<br>(N=34) | Control<br>(N=26) |
| --- | --- | --- | --- |
| Age at diagnosis/enrollment in years (mean) | 46.87 | 57.58 | 42.30 |
| Sex (M/F) | (28/55) | (9/25) | (8/18) |
| Race | White | white | White## |
| BMI kg/m<sup>2</sup> (mean ) | 30.35 | 28.33 | 27.64 |
| EBV flag *** | 5 | 0 | 0 |
| **Therapy** |  |  |  |
| Disease modifying therapy | 77 | 0 | 0 |
| #Combination Therapy | 6 |  |  |

\*Group 1= Treated RRMS patients
\*\* Group 3= Treatment naïve RRMS patients
\*\*\*EBV flag=Epstein-Barr virus previous infection

**Table 2:**
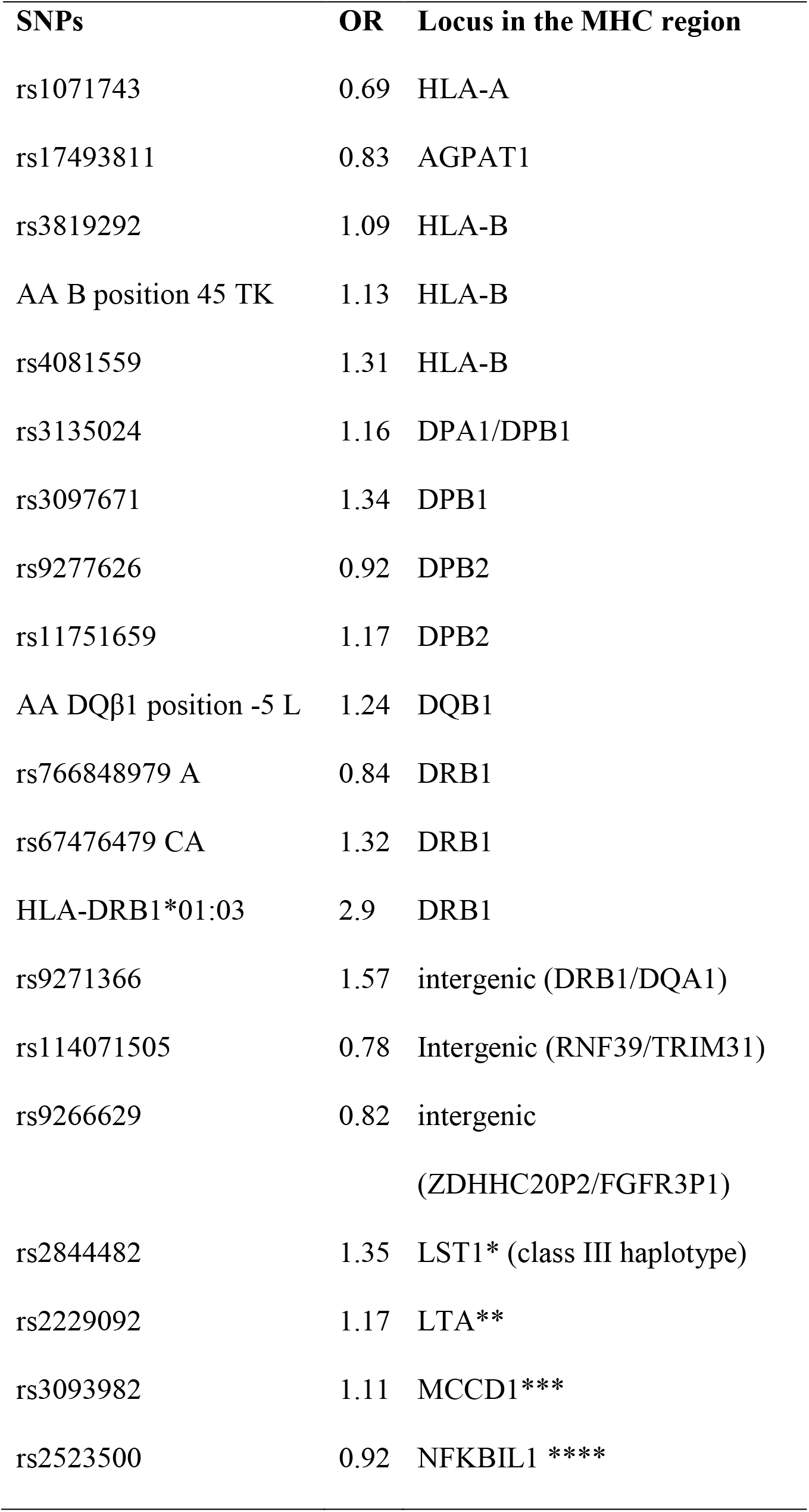
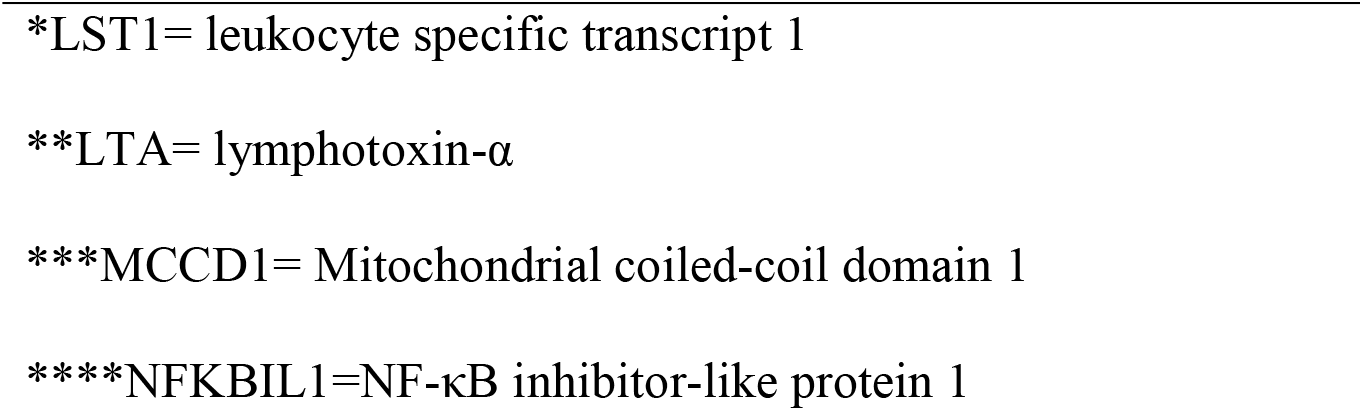
Studied SNPs associated with MS inside MHC region.

| **SNPs** | **OR** | **Locus in the MHC region** |
| --- | --- | --- |
| rs1071743 | 0.69 | HLA-A |
| rs17493811 | 0.83 | AGPAT1 |
| rs3819292 | 1.09 | HLA-B |
| AA B position 45 TK | 1.13 | HLA-B |
| rs4081559 | 1.31 | HLA-B |
| rs3135024 | 1.16 | DPA1/DPB1 |
| rs3097671 | 1.34 | DPB1 |
| rs9277626 | 0.92 | DPB2 |
| rs11751659 | 1.17 | DPB2 |
| AA DQβ1 position -5 L | 1.24 | DQB1 |
| rs766848979 A | 0.84 | DRB1 |
| rs67476479 CA | 1.32 | DRB1 |
| HLA-DRB1*01:03 | 2.9 | DRB1 |
| rs9271366 | 1.57 | intergenic (DRB1/DQA1) |
| rs114071505 | 0.78 | Intergenic (RNF39/TRIM31) |
| rs9266629 | 0.82 | intergenic<br>(ZDHHC20P2/FGFR3P1) |
| rs2844482 | 1.35 | LST1* (class III haplotype) |
| rs2229092 | 1.17 | LTA** |
| rs3093982 | 1.11 | MCCD1*** |
| rs2523500 | 0.92 | NFKBIL1 **** |

\*LST1= leukocyte specific transcript 1
\*\*LTA= lymphotoxin- $\alpha$
\*\*\*MCCD1= Mitochondrial coiled-coil domain 1
\*\*\*\*NFKBIL1=NF- $\kappa$ B inhibitor-like protein 1

**Table 3:** Comparative relative genera abundance between Cluster A and Cluster B.

| **Genus** | **Cluster A** | **Cluster B** |
| --- | --- | --- |
| *Blautia* | 18.06% | 16.50% |
| *Coprococcus* | 9.28% | 8.89% |
| *Akkermansia* | 8.92% | 5.51% |
| *Bacteroides* | 6.58% | 5.73% |
| *Ruminococcus* | 9.38% | 9.82% |
| *Bifidobacterium* | 4.10% | 6.17% |
| *Collinsella* | 2.95% | 4.41% |
| *Streptococcus* | 3.07% | 2.72% |
| *Roseburia* | 2.75% | 1.02% |
| *Faecalibacterium* | 3.33% | 2.06% |
| *Methanobrevibacter* | 2.41% | 2.12% |
| *Dorea* | 2.33% | 3.28% |
| *Prevotella* | 1.07% | 1.08% |
| *Alistipes* | 1.00% | 1.09% |
| *Clostridium* | 0% | 1.52% |
| *Oscillospira* | 1.41% | 1.61% |
| *Phascolarctobacterium* | 1.05% | 0 |
| *Megasphaera* | 0 | 1.52% |

## Results

### Demographics and summary of electronic health record from the study participants

The number of patients in Group 1 and 3 were 83 and 34 respectively. The average age of MS patients in Group 1 and Group 3 at diagnosis were 46.87 and 57.58 years respectively. Their BMIs were 30.35 and 28.33 for Group 1 and Group 3, respectively. Seventy seven patients in Group 1 were on a single DMT while six patients were on two different DMTs in the last 6 months of the time of fecal samples collection (Table 1).

### The gut microbiome profile of Group 1 (Treated RRMS) and Group 3 (Treatment naïve RRMS) MS cases and controls

The Faith’s phylogenetic diversity between the MS cases and non-MS healthy control was significantly different (Figure 2A, *P* value = 0.002) and so was the Pielou’s evenness index (Figure 2B, *P* value = 0.03). However, the Shannon diversity index between the cases and controls was not significantly different (Figure 2C). When we compared the Faith’s phylogenetic diversity between Group 1, Group 3, and healthy control group (see materials and methods), we observed that while both Group 1 and Group 3 were significantly different from the healthy control group, the difference was not significant between the two case groups (Figure 3).

**Figure 2.**
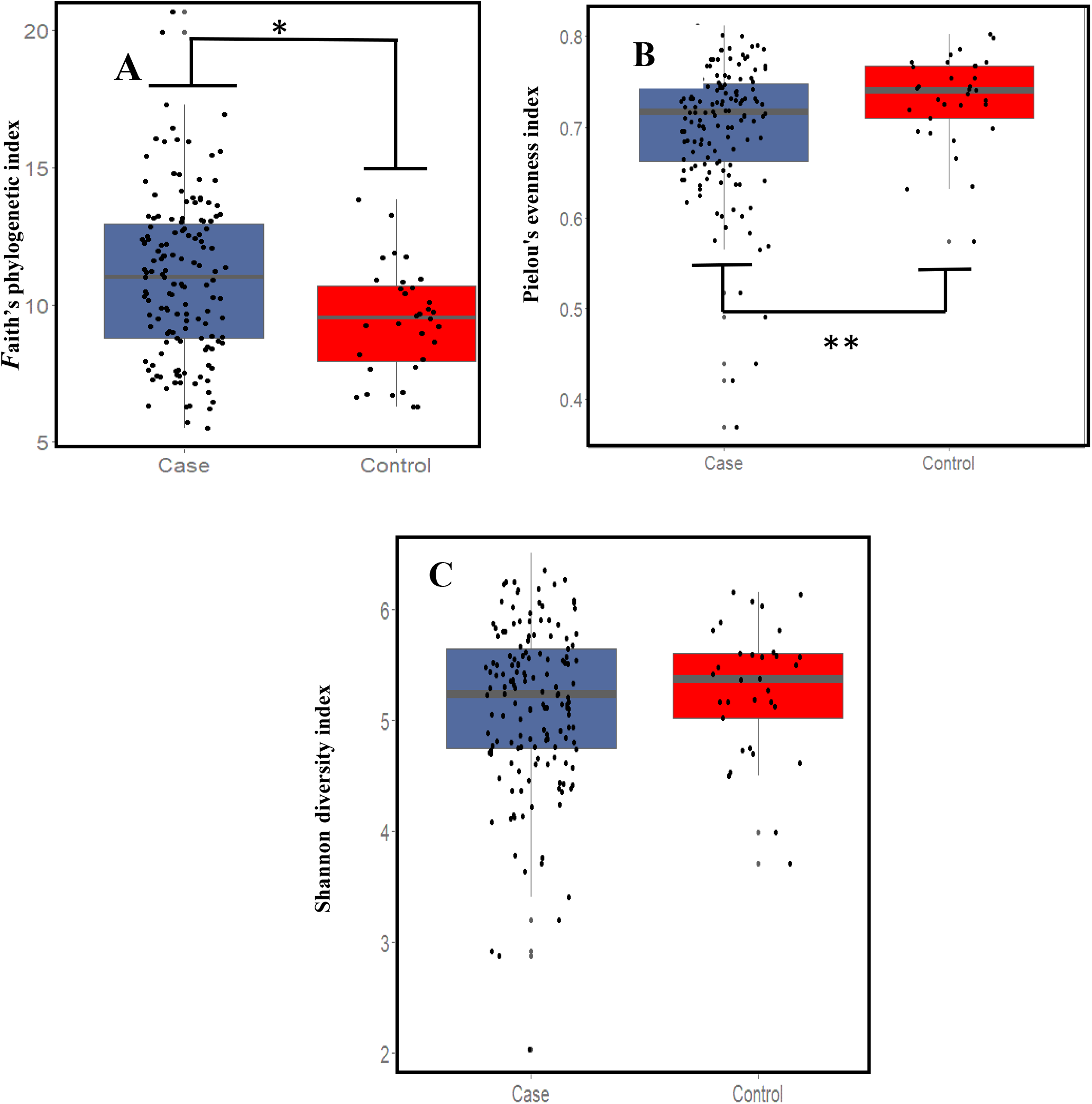
A. The boxplot representing Faith’s Phylogenetic diversity (PD) where there was a significant difference between MS cases and control. B. Pielou’s evenness index where there were a significant difference between cases and controls. C. Shannon-wiener diversity index (H) where both MS cases and control microbiome were similar, Kruskal Wallis test was used to detect any significant differences between cases and controls in different indices of alpha diversity

**Figure 3.**
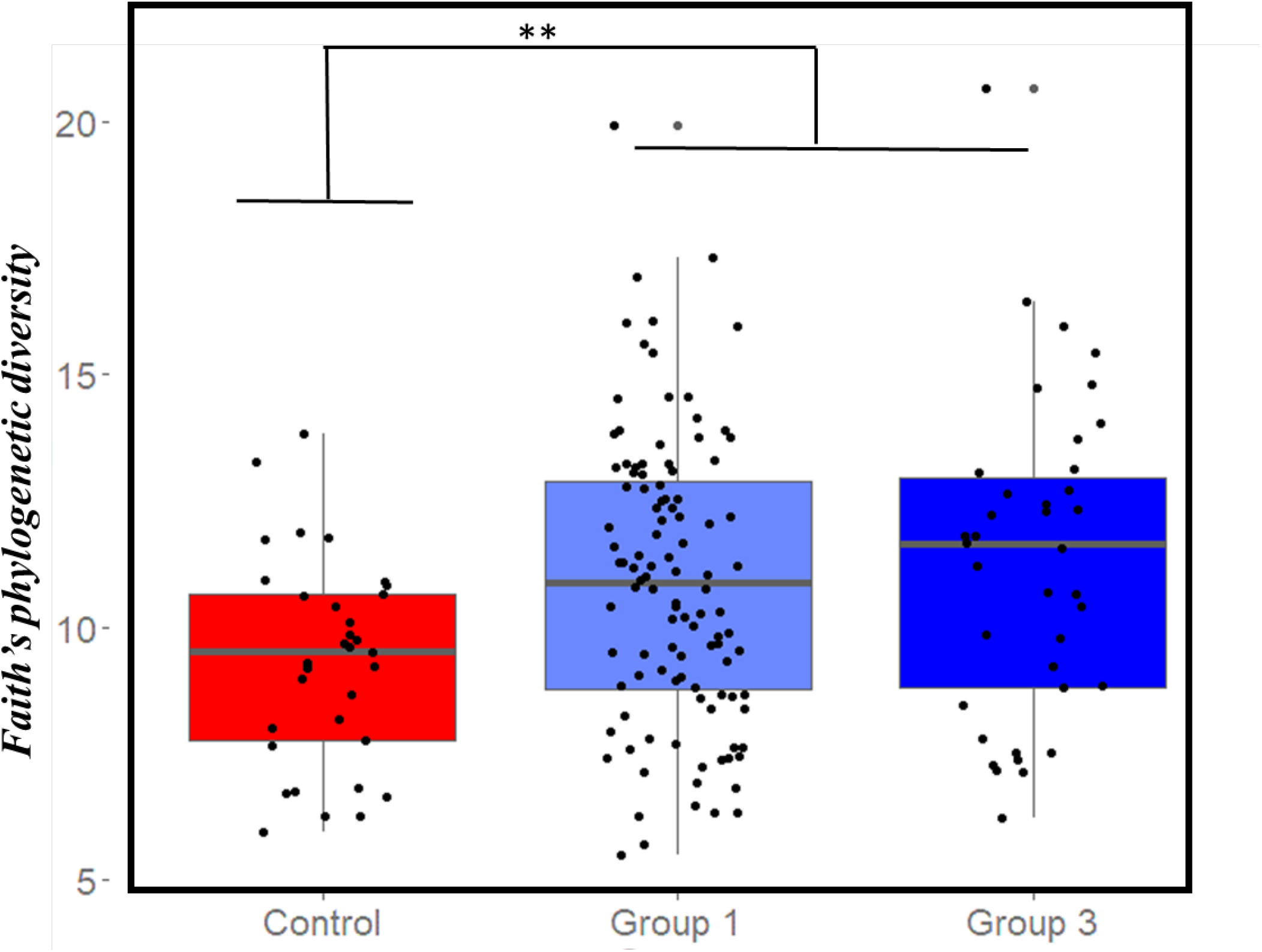
Box plot representing the Faith’s phylogenetic diversity in MS patients group 1 (Group 1, treated RRMS), group 3 (Group 3, treatment naïve RRMS), and healthy controls. There was a significant difference between the control and each group of MS cases individually. On the other side, there was no significant difference between the two groups of MS cases.

### Identification of a unique MS patients cluster

When we performed the unweighted UniFrac principal component analysis (PCA) on microbiota of 117 cases and 26 controls, we observed two clusters, a large cluster named Cluster A consisting of 98 cases and 26 control (n=124) and a smaller cluster named Cluster B consisting of 19 cases only (Figure 4). PC1 accounted for 15.1% of the variation, while PC2 accounted for 9.94% of the variation. These two clusters were significantly different by the Permanova test (*p* value=0.01). However, differences in these two clusters were not associated with age, DMT used, number of MRI lesions or any other disease conditions like gastric issues.

**Figure 4.**
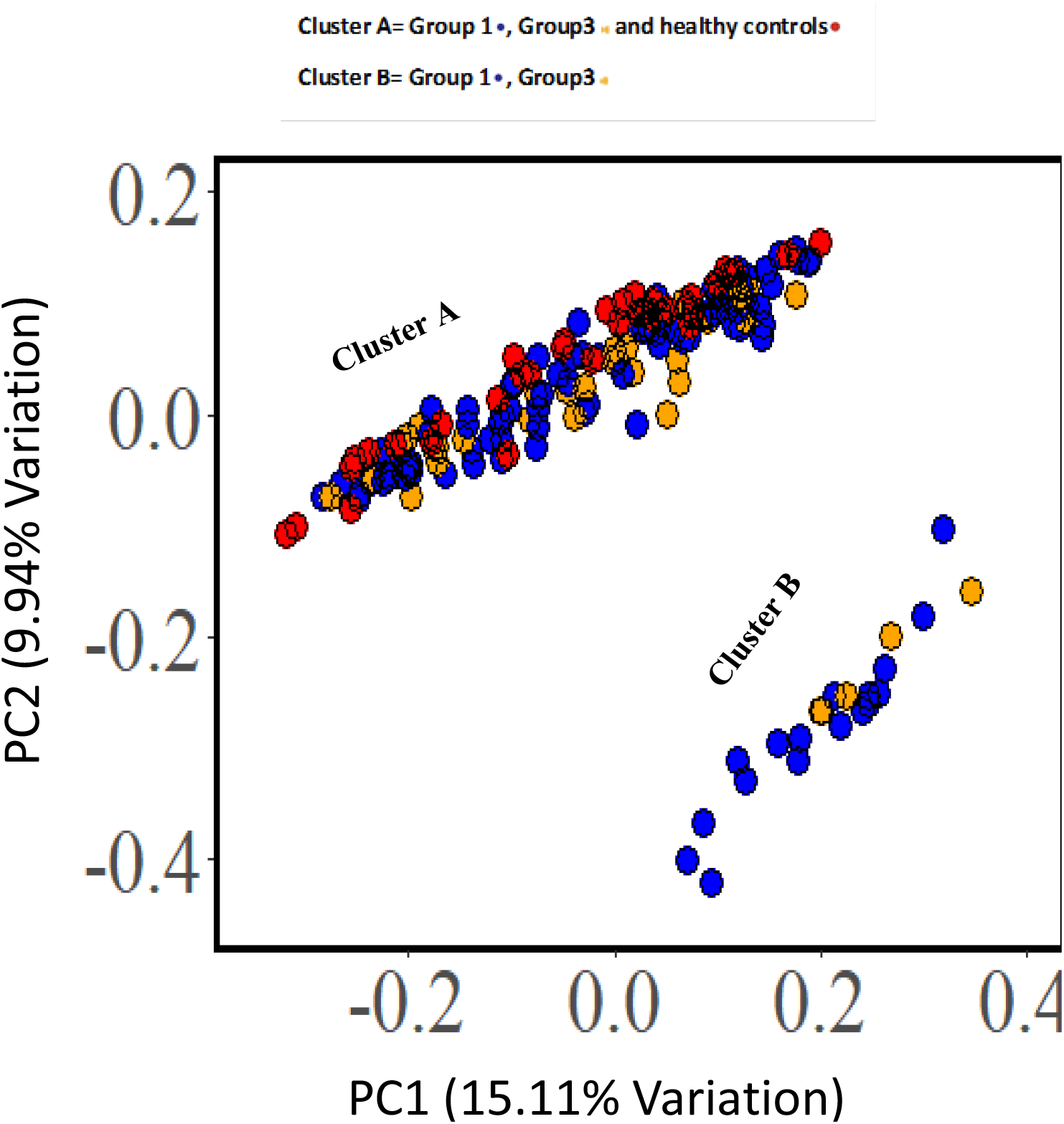
Unweighted UniFrac Principal Coordinate (PCA) of groups one (treated RRMS), three (treatment naïve RRMS) and healthy controls showed two distinct clusters (A and B). Each dot represents a MS case or healthy control and the PCA plot show the abundant taxa in each patient gut microbiota. The two chosen PC coordinates showed the most diversity, represented in percentage on the axis.

### Relative abundance analysis

We observed several differences in relative abundances of different phyla at 99% cutoff, but they were not statistically significant. For example, *Actinobacteria* showed a higher relative abundance in Cluster B whereas *Verrucomicrobia* showed a higher relative abundance in cluster A (Figure 5A, and Supplemental Table 2). Both *Bacteroidetes* and *Firmicutes* showed comparable abundances in the two clusters. Interestingly, *Proteobacteria* was not detected in cluster A. As shown in Figure 5B and ST3, *Lachnospiraceae* family showed small difference between the two clusters. At the genus levels, there were some genera such as *Phascolarctobacterium* and *Clostridium* and *Megasphaera* were not detected in Cluster B and in Cluster A respectively (Figure 5C and Supplemental Table 3). Moreover, *Bifidobacterium* showed higher abundance in Cluster B while *Akkermansia* in cluster A.

**Figure 5.**
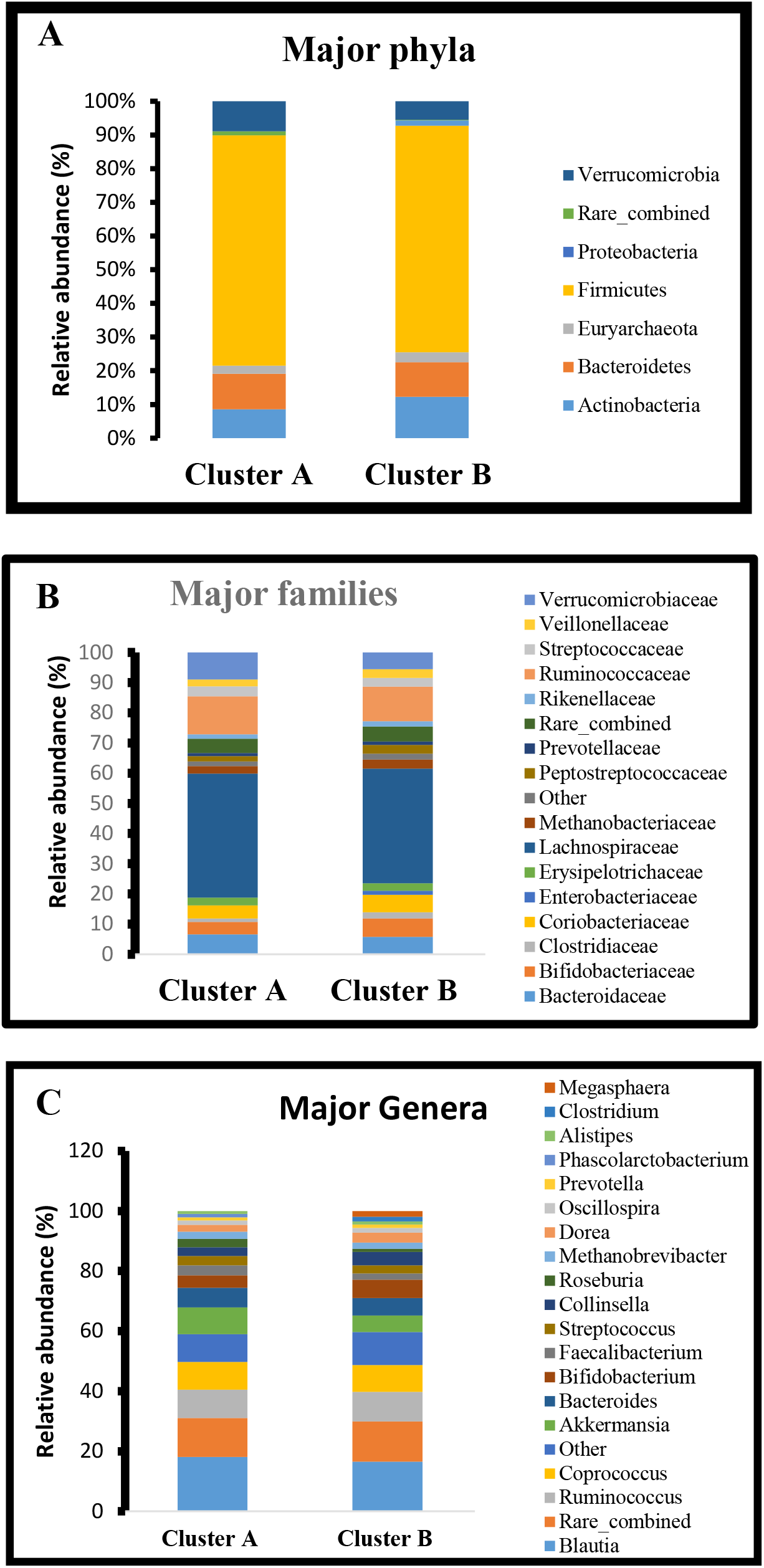
Relative abundance of major phyla (A), major families (B), and major genera (C) in the clusters A and B generated from the Unweighed Unifrac

### The QCAT-GEE tests showed difference between Cluster A and Cluster B

After applying all three tests of QCAT-GEE to the taxonomy table with all ranks from kingdom to genus, we observed that QCAT-GEE two-parts test and the positive test showed that only *Porphyromonadaceae* family was significantly different between two clusters A and B (p-value is 0.00899).

### Polygenic risk scores associated with Clusters A and B

Since MS has a strong genetic susceptibility component, we used a validated risk score method with MS disease [10,40] in our study. The healthy control subjects tended to have a lower polygenic risk score (from 0.007 to 0.017) whereas, the MS cases tended to have a higher polygenic risk score from 0.007 to 0.022 (Figure 6). The t-test also showed high significant differences between the polygenic risk scores between cases and controls (p value= 2.682e-05). When considering both the microbiome diversity based clusters and the polygenic risk scores together (Figure 7), an interesting trend was observed where the gut-microbiome of subjects of Cluster A, which included a significant number of both cases and controls, tended to have a lower polygenic risk scores compared to cluster B (higher polygenic risk score) which consisted of cases only, and it was also statistically significant by ANOVA test (p value= 1.56 *10^−5^). This suggests that the patients’ with higher polygenic risk score may be associated with their gut microbiome. Additionally, the cases with lower genetic risk scores tend to have their microbiome closer to the healthy controls compared to the cases with a higher genetic risk score.

**Figure 6.**
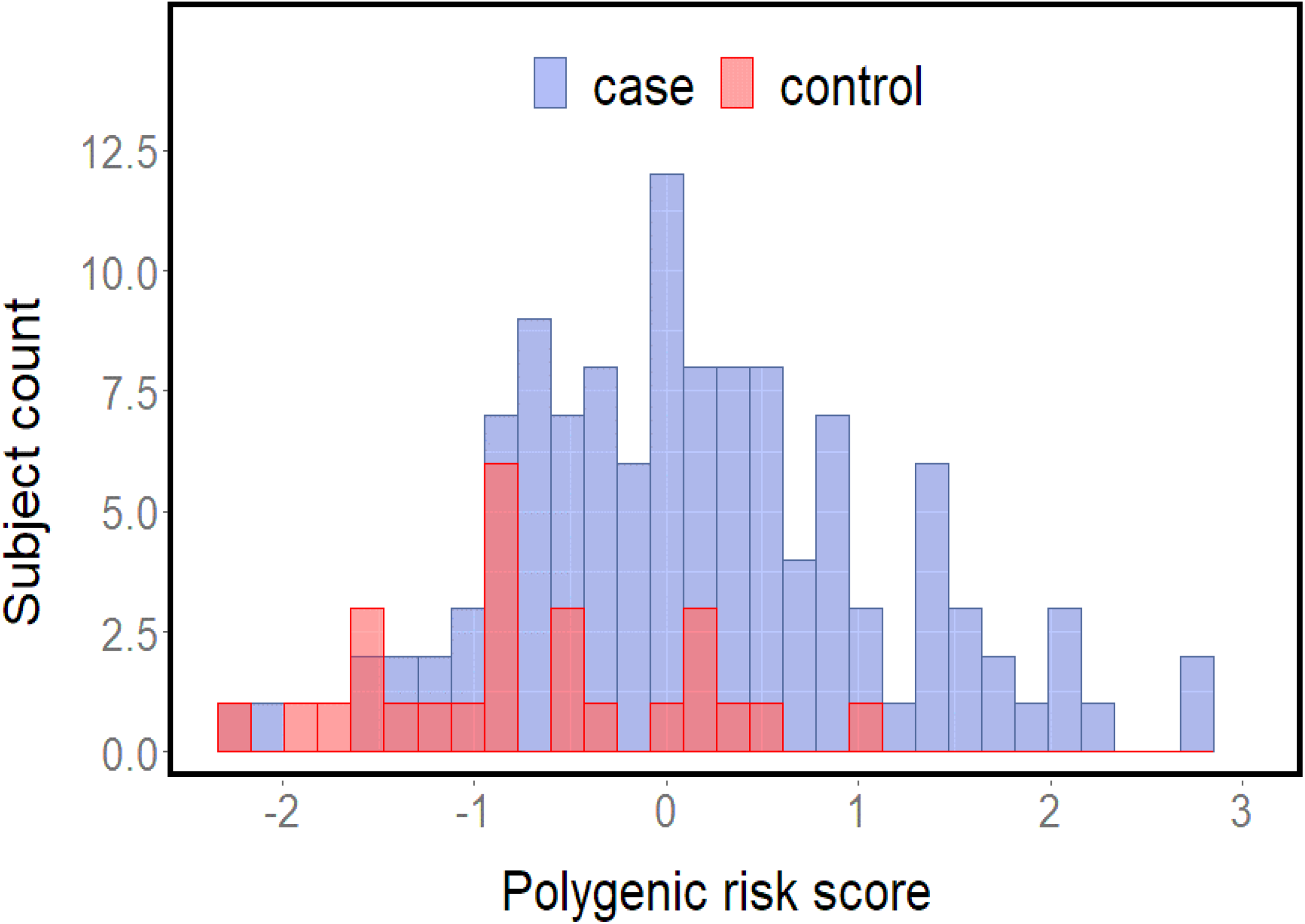
A Histogram showing the distribution of polygenic risk score in our cohort of both healthy controls (red bars) and MS cases (blue bars). Here, the histogram showed the controls having low genetic risk scores in comparison to the MS cases. The polygenic risk scores were tested using SNPs both inside MHC region (Table 2) and outside MHC region (supplementary table 1). T-test showed high significant difference between cases and control (p value= 2.682e-05).

**Figure 7:**
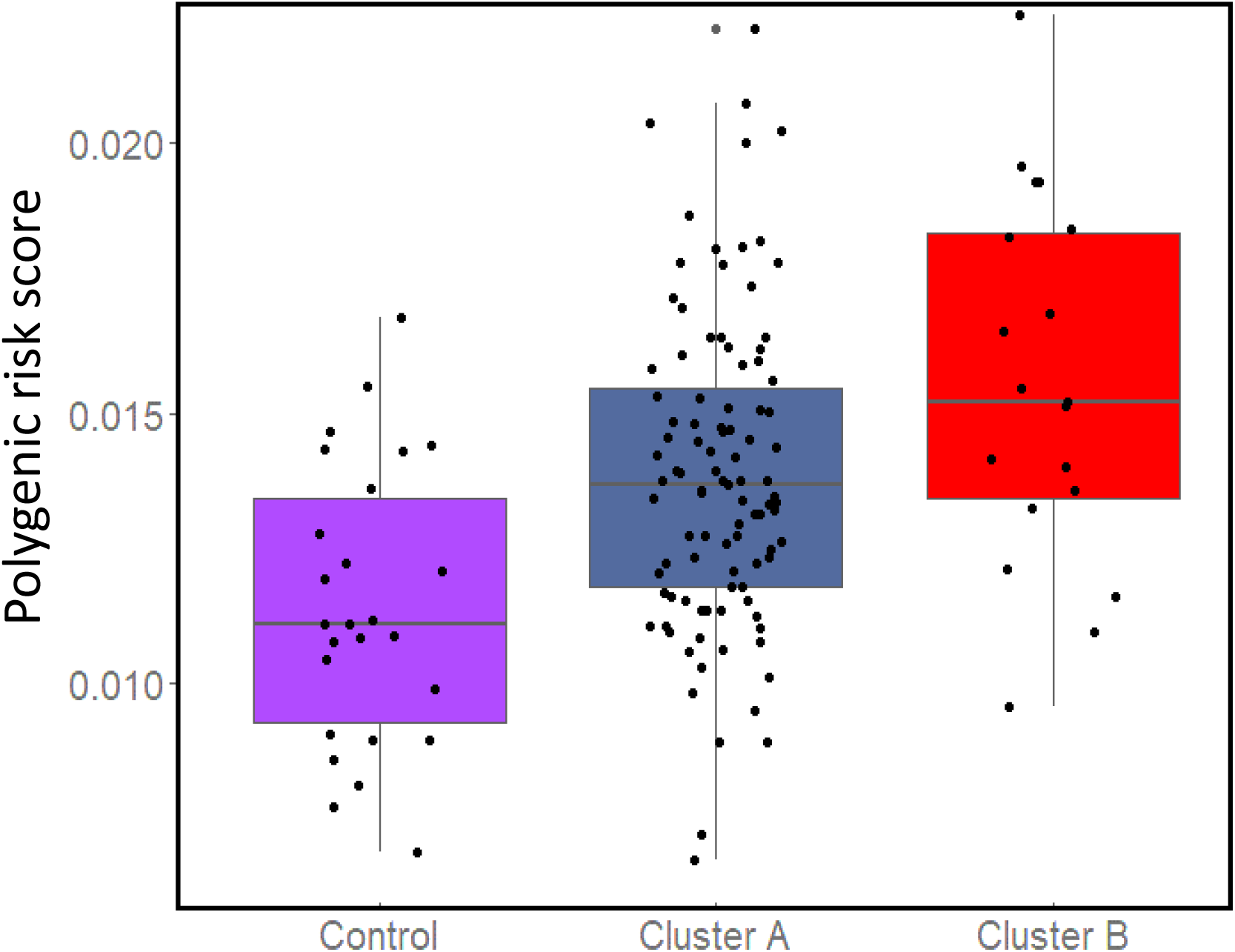
Comparison of polygenic risk score of the MS cases in Cluster A and Cluster B with that healthy controls from the Unweighted unifrac analysis. This plot showed the rising of polygenic risk scores from low values in healthy controls to higher values in the both clusters comprising of MS cases. However, the cluster B showed the highest genetic risk score. ANOVA test to check difference in variances between these three groups showed significant difference (p =1.56e-05).

## Discussion

Unravelling the genetic-environmental factor(s) that control susceptibility to a complex autoimmune diseases such as MS is challenging. Studies have identified the genetic susceptibilities to MS [10,12,13] and association of individual microbes and/or gut dysbiosis in MS [5,6,16]. In this study, we calculated a polygenic risk score of MS patients in our cohorts and conducted gut microbiome analyses to see the association between high-risk cohorts and identify unique microbiota signatures in their gut. We calculated the genetic risk score based on a validated risk scores for the MS disease as described before [10]. Since MS is a complex disease, many variants share the responsibility in increasing the patients’ susceptibility to this devastating disease. Surprisingly, we found that patients exhibiting the highest genetic risk score are the patients who had a distinct microbiome.

Through studying the demographics data of the patients enrolled in this study and their Epstein-Barr Virus (EBV) status in the electronic health record, we made several observations. Notably, only five out of the tested MS patients had evidence of EBV infection in their electronic health records (see table 1). This was surprising as EBV was recently described as a virus that increases the risk for MS susceptibility [41]. As expected, the female ratio was higher than males which is known in MS disease epidemiology [42].

We observed a significant difference in both Faith’s phylogenetic diversity and Pielou’s evenness indices between the MS cases and healthy controls. The higher significant faith phylogenetic diversity MS cases was in agreement with the Unweighed Unifrac analysis. Absence of any significant differences in Shannon index between our MS case and healthy control cohorts was similar to a couple of previously published results, especially if the gut microbiome was collected during non-active episodes of the MS disease [4,43]. Indeed, Chen et al (2016) reported that patients with active episodes of RRMS have a decline in the species richness [16]. There were no significant differences in alpha and beta diversity indices patients treated with DMTs were compared with treatment naïve patients, Some studies have reported changes in gut microbiome composition after treatment especially with glatiramer acetate (GA) and dimethyl fumarate (DMF) [20,44]. Future studies should consider assessing the gut microbiome of MS patients at different time points in RRMS to ascertain different dysbiosis state during the active and non-active phases of the disease. Surprisingly, we observed a significant number of MS patients who had a higher polygenic risk score than the healthy controls but had a similar gut microbiome compared to the controls. We speculate that this could be due to fact that i) 94% (91 out of 98) of the patients were in remitting phase of which 68% (67 out of 98) were also on one or more DMTs. The Unweighed unifrac analysis is sensitive to rare lineages within a microbial community [47,48]. Based on our QCAT analysis,*Porphyromonadaceae*, a typically low abundance family, was observed to be significantly higher in Cluster B. This family was also found to be associated with worse EAE outcomes in genetically susceptible mice [49]. Moreover, its presence has been correlated with IL17, an interleukin known to be upregulated in MS which is also high in the ileum in other metabolic diseases like obesity and diabetes [50]. Even in neurodegenerative diseases such as ankylosing arthritis [51] and Parkinson diseases, *Porphyromonadaceae* was enriched. [52]. Lower number of patient samples in Cluster B limited the identification of other significant taxa associated with high genetic risk score patients. Meanwhile, another species that showed higher abundance in Cluster A without showing significance was *Akkermansia* as reported in Mirza et al (2020) [53].

Genetic studies conducted to detect the variants causing MS disease reported the modest effect to variant HLA DRB1*15:01 and many other loci with smaller associations [54]. However, no study has pointed the risk of MS disease to a certain allele in the HLA class II because all of the alleles in this region, especially, in European ancestors are inherited together due to intense linkage disequilibrium [56]. Moreover, some of these detected SNPs for MS risk in the literature are common and can be present in healthy unaffected individuals [55]. Thus, polygenic risk score measurement is suitable in this complex disease to reveal the cumulative risk to MS. Good predictability has been achieved by measuring this score before in other diseases (prostate cancer and systemic lupus erythematosus) including MS disease [57–59]. In our cohort, as expected, the MS patients have higher polygenic risk score than healthy controls. However, we found a unique cohort with the highest risk score having a unique gut microbiome. It is not clear at this stage, whether the dysbiotic gut microbiome increased the MS risk together with the genetic susceptibility or the host genotype affected the gut microbiome composition. For instance some studies have suggested the heritability of the gut microbiome [5]. Studies have also suggested that host genes affect the shape of the gut habitat thereby leading to variation in the gut microbiome [60]. Furthermore, in case of the MS, variants in MHC region in general could affect the shaping the gut microbiome through restricted colonization of some bacterial species through either their immune elimination or their inability to adhere to the intestinal epithelium [61] through the IgA mediated selection [62]. In addition, MHC region affects the T-cells maturation which subsequently can affect its autoreactivty [63]. Indeed, all of these studies support the interaction of genes and gut microbiome in precipitating different diseases.

## Conclusion

In summary, we showed that a small cohort of MS patients showed high polygenic risk score who also harbored a distinct microbiota in the gut. This observation droves that idea that indeed, both genetic susceptibility and environmental factor, dysbiosis of the gut microbiome is associated with MS, albeit in a small number of patients. While future studies with larger sets of patients are needed to confirm the relationship between the genetic risk score and the MS gut microbiome, we believe our study provides a foundation for such a study.

## Authors’ contributions

SKS, PA, KA, and RV conceived and designed the study, TK along with SKS and PA enrolled the study participants, NE and TL performed the laboratory work, SKS, VG, NE, RV, VB, QL, JM, ZT, TL, and TK analyzed the data, NE, SKS, RV, wrote the first draft and all other authors subsequently contributed to the writing and editing of the manuscript.

## Acknowledgements

We would to thank all study participants including MS patients who volunteered their time and graciously provided biological samples. Without their support, this study could not have been completed.

